# COL7A1 expression improves prognosis prediction for patients with clear cell renal cell carcinoma atop of stage

**DOI:** 10.1101/2023.01.16.524247

**Authors:** Dzenis Koca, Irinka Séraudie, Rémy Jardiller, Claude Cochet, Odile Filhol, Laurent Guyon

## Abstract

Clear cell renal cell carcinoma (ccRCC) accounts for 75% of kidney cancers. Due to the high recurrence rate, and treatment options that come with high costs and potential side effects correct prognosis of patient survival is essential for the successful and effective treatment of patients. Novel biomarkers could play an important role in the assessment of the overall survival of patients. COL7A1 encodes for collagen type VII, a constituent of the basal membrane. COL7A1 is associated with survival in many cancers; however, the prognostic value of COL7A1 expression as a standalone biomarker in ccRCC has not been investigated. We used Kaplan-Meier curves and Cox proportional hazards model to investigate the prognostic value of COL7A1, as well as Gene Set Enrichment Analysis to investigate genes that are co-expressed with COL7A1. COL7A1 expression was used to stratify patients into four groups of expression, where the 5-year survival probability of each group was 72.4%, 59.1%, 34.15%, and 8.6% in order of increasing expression. Additionally, COL7A1 expression was successfully used to further divide patients of each stage and histological grade into groups of high and low risk. Similar results were obtained in independent cohorts. *In-vitro* knockdown of COL7A1 expression significantly impacted ccRCC cells’ ability to migrate and proliferate. To conclude, we identified COL7A1 as a new prognosis marker that can stratify ccRCC patients.

## 1. Introduction

In the year 2020, around the world, there was an estimated 431,300 new cases of kidney cancer, accounting for 2.2% of all cancer cases, and nearly 179,300 patients died of this disease [1]. Clear cell renal cell carcinoma (ccRCC) is a distinct histologic subtype of kidney cancer that accounts for approximately 75% of kidney cancer cases [2]. ccRCC is a highly metastatic cancer; nearly 17% of patients have developed metastasis at the time of diagnosis [3]. The ccRCC subtype is histologically characterized by having transparent (clear), round-shaped cells. The reason for discoloration is the accumulation of lipids inside tumor cells, as the cell goes through epithelial to mesenchymal transition (EMT) and partial adipogenic trans-differentiation [4]. Multiple cancer driver events were identified through the molecular characterization of ccRCC. Some of the most common driver events include mutations and methylation of VHL, PBRM1, BAP1 and SETD2 genes, as well as changes in a chromosomal structure such as loss of 3p and gain of 5q chromosomes [5]. Over the years, the prognosis of ccRCC patients has improved, although its incidence is indeed increasing[3]. According to the 2019 data, around 17% of newly diagnosed patients have developed distant metastasis, and the 5-year survival probability of these patients is less than 12% [3]. Early ccRCC cases are asymptomatic and they are often discovered accidentally, during standard medical checkups and CT or ultrasound of the abdomen. Still, reoccurrence occurs in around 30% of patients that undergo surgery, and this includes 10-25% of patients with localized disease [6]. Novel therapeutical options such as targeted therapies and immuno-therapies often come with side effects and high costs and are often prescribed to patients with intermediate to poor prognoses [7]. Nevertheless, there is an urgent need for biomarker-driven studies with an overarching goal to incorporate newly proven biomarkers for innovative trial designs and to give a more accurate and individualized prognosis to patients.

Extracellular matrix (ECM) plays an important role in many cancers. ECM can serve both as mechanical support/scaffold for tumor growth and cell proliferation and migration as well as a modulator of biochemical cues by promoting epithelial-mesenchymal transition, sustaining self-renewal, inducing metabolic reprogramming and accumulating and delivering various growth factors [8]. Collagen proteins are major structural and functional components of ECM. The basement membrane Type I collagen is a prevalent component of the stromal ECM, its expression is spatially and temporally regulated to maintain normal cell behavior. Type VII collagen is a protein that belongs to the family of collagens, which is encoded by the Collagen Type VII Alpha 1 Chain (COL7A1) gene. Collagen 7 fibril is formed by intertwining three identical collagen chains, and it primarily functions as an anchoring fibril between the basal membrane and proximal cells of stratified squamous epithelia [9]. Germline mutations in COL7A1 are one of the main causes of dystrophic epidermolysis bullosa, which in turn can increase the chance of squamous cell carcinoma [10]. Increased expression of COL7A1 was also detected in highly metastatic cancer cell lines, as well as in prostate cancer-initiating cell spheroids [11,12]. COL7A1 expression has already been investigated as a potential prognostic biomarker in patients with lung squamous cell carcinoma [13], esophageal squamous cell carcinoma [14] as well as gastric cancer [15].

Assessment of patient survival could improve decision-making in a clinical environment. Additionally, certain treatment options are allowed to be used only in cases of poor prognosis. Clinical and biochemistry markers, such as stage and grade of tumor, are often used in the assessment of ccRCC prognosis [16]. Recently, we have shown that, for most cancers, survival prediction can be improved by incorporating clinical and transcriptomic data [17]. Additionally, there is an increasing interest in how ECM-related genes could be used to assess the survival of ccRCC patients [18–21]. To the best of our knowledge, COL7A1 expression in ccRCC was not investigated as an independent prognostic biomarker in this disease.

The present paper is organized as follows: Firstly, we performed a screening for potential biomarker genes in ccRCC followed by an in-depth investigation of the top results. Secondly, we investigated the possible biological implications of COL7A1 through the assessment of co-expressed genes. Finally, we perform in vitro experiments to investigate the impact of COL7A1 expression on the proliferation and migration of ccRCC cell lines.

## 2. Materials and Methods

### Datasets and processing

The Cancer Genome Atlas (TCGA) is a cancer genomics program that characterized over 20.000 tumoral and peritumoral tissue samples of 33 different cancers, including ccRCC (with the acronym KIRC). The transcriptome profiling data of primary tumor tissue and corresponding clinical and pathological information of TCGA:KIRC cohort were obtained using the “TCGAbiolinks” R package (v2.62)[22]. As per a recent suggestion [23], we downloaded harmonized transcriptomic data, which was already Transcripts Per Million (TPM) normalized. To decrease computational time, only the protein-coding genes, with total expression across samples greater than 10 TPM were retained. Data was further transformed in *log*2(TPM+1) counts. To deal with multiple samples per patient, only samples that could be found in GDS “FireBrowse” (http://firebrowse.org/, accessed on 2 November 2022) were used in downstream analysis.

To validate the results on independent cohorts, we have obtained three additional datasets. Transcriptome profiling data of the E-MTAB-1980 dataset was obtained from Array express, while clinical data was obtained from supplementary data of the original publication [24]. The GSE167093 dataset was obtained from Genome Expression Omnibus (GEO)[25]. “Braun *et al*.” dataset was obtained from the supplementary data of the corresponding publication [26]. All datasets of independent cohorts were already normalized by the original authors. Details on normalization are available in respective publications. An overview of clinicopathological characteristics of patients in datasets, for which clinical data was publicly available, is available in Table 1.

**Table 1.** Clinicopathological characteristics of patients with ccRCC in four different datasets. Presented are datasets that had publicly available clinicopathological data.

|  | TCGA | E-MTAB-1980 | GSE167093 | Braun *et al.* |
| --- | --- | --- | --- | --- |
| **Total samples** | 532 | 101 | 604 | 225 |
| **Median age at diagnosis years (range)** | 61 (26-90) | 64 (35-91) | 62 (23-85) | 62 (30-88) |
| **Sex, N (%)** |  |  |  |  |
| Female | 187 (35.15) | 24 (23.8) | 247 (41) | 61 (27.1) |
| Male | 345 (64.85) | 77 (76.2) | 357 (59) | 164 (72.9) |
| **Tumor grade<sup>a</sup>, N (%)** |  |  |  |  |
| I | 14 (2.7) | 13 (13.1) | 100 (18.8) | - |
| II | 228 (43.5) | 59 (59.6) | 304 (57) | - |
| III | 206 (39.3) | 22 (22.2) | 105 (19.7) | - |
| IV | 76 (14.5) | 5 (5.1) | 24 (4.5) | - |
| Missing | 8 | 2 | 71 | 225 |
| **Tumor stage<sup>a</sup>, N (%)** |  |  |  |  |
| I | 266 (50.3) | 66 (65.3) | 306 (50.7) | 0 |
| II | 57 (10.8) | 20 (19.8) | 98 (16.2) | 0 |
| III | 123 (23.2) | 3 (3) | 138 (22.8) | 0 |
| IV | 83 (15.7) | 12 (11.9) | 62 (10.3) | 225 (100) |
| Missing | 3 | 0 | 0 | 0 |
| **EVENTS** | 175 (32.9) | 23 (22.78) | - | 173 (76.9) |
| **5 year survival probability % (95% CI)** | 62.9 (58.2-68.1) | 79.6 (71.7-88.3) | - | 20.4 (15.5-26.9) |

a) Samples with missing information were not included into percentage calculation

### Univariable and multivariable Cox model

Cox proportional hazards model (“Cox model” in the rest of the text) is one of the most popular methods used in survival analysis. The advantage of Cox model is that it is applicable on continuous variables (without the need to set a threshold) and that it can deal with the right censored data. Cox model can be represented as:

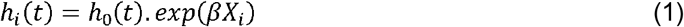

where *h(t)* is the instantaneous risk of dying for patient *i = 1,…,n, h*_*0*_*(t)* a common hazard baseline for all patients, β a coefficient to be estimated, and *X*_*i*_ the expression of a given gene in the tumor sample of patient *I* [27]. If β is significantly positive (resp. negative), the risk increases (resp. decreases) with gene expression: the gene could act as an oncogene (resp. a tumor suppressor). In case of multivariable Cox model, β*X* from equation 1 is transformed to ∑β_*j*_*X*_*j*_, with *X*_*j*_ being one of the variables, and β_*j*_ being a learned coefficient for variable *X*_*j*_.

In order to select genes which could be used in a clinical environment, we applied two thresholds on genes expression: a) filtering out genes with extremely low expression (median expression = 0 in log2(TPM+1)), b) filtering out genes that are not variable enough to be precisely measured in a clinical environment (quantile 80 – quantile 20 = 0, in log2(TPM+1)). After the filtering process, we used each remaining gene to train a univariable Cox model. For this purpose, we used the R “survival” package (v3.3.1) [28] and we considered the obtained P-values to judge the goodness of fit of the Cox model.

Additionally, we performed a univariable Cox model on clinical covariates such as the age of a patient, gender, clinical stage, and histological grade, as well as COL7A1 expression. We then used all mentioned covariates to learn a multivariable Cox model, and compared changes in a P-value of COL7A1 covariate between the univariate and multivariate Cox model.

We conducted an analysis of deviance of the multivariable Cox model to determine the added value of COL7A1 expression, compared to the multivariable Cox model based solely on clinical covariates alone. The analysis of deviance table presents the change in a predictive power of model as each of the covariates is added to the model, while the P-value shows whether the improvement was significant.

### Kaplan-Meier curves

Patients were divided into groups of High and Low COL7A1 expression using a cut-point determined by the “surv_cutpoint” function of the “survminer” package (v0.4.9) [29]. We applied the same process in the other two datasets for which overall survival information was available. Division of patients according to COL7A1 expression in five groups was performed by dividing the outlier-free range of expression into five equal sections using the “cut” function. We computed and drew all Kaplan-Meier curves using packages “survival” and “survminer”.

### GSEA

To better understand the biological implication of COL7A1 aberrant expression, we investigated genes that are co-expressed with COL7A1. Gene set enrichment analysis (GSEA) allows us to investigate how does the expression of a set of genes that belongs to a common biological pathway, varies in expression together with COL7A1 [30]. We performed GSEA using the “ClusterProfiler” (v4.6.0) package [31]. One of the requirements for GSEA is an ordered vector of genes, based on which enrichment will be calculated. For this purpose, we investigated co-expression through Pearson’s correlation between COL7A1 expression and the expression of other genes available in each investigated dataset individually. Gene sets (pathways) on which we have performed GSEA were obtained from the Molecular Signature Database (MSigDB, v2022.1), with a focus on the Hallmark dataset [30,32].

### Statistical analysis and data manipulation

All of the aforementioned analyses were performed using the R software (v4.2.1)[33]. The set of packages contained in the “tidyverse”(v1.3.2) library (dplyr, ggplot, purr…) were used to manipulate data and create graphs [34].

### Cell culture

RCC cell lines 786-O, ACHN and Caki-1 were obtained from ATCC (CRL-1932, CRL1611 and HTB-46, respectively). The cell lines were grown in 10 cm diameter plates in a humidified incubator (37°C, 5% CO_2_) with RPMI 1640 medium (Gibco) containing 10% of fetal bovine calf serum, penicillin (100 U/mL) and streptomycin (100 μg/mL).

### Generation of shCOL7A1 786-O cells

Four Lenti-pLKO.1-puro shRNA vectors that specifically targeted COL7A1 mRNA sequences were chemically synthesized (Sigma Aldrich) and included in viral particles:

(#1: 5′- CCGGCCCTTGAGAGGTGACATATTCCTCGAGGAATATGTCACCTCTCAAGGGTTTTTG-3′, #2: 5’- CCGGGCTCGCACTGACGCTTCTGTTCTCGAGAACAGAAGCGTCAGTGCGAGCTTTTTG-3′, #3: 5′- CCGGGAGCCAGTGGATTTCGGATTACTCGAGTAATCCGAAATCCACTGGCTCTTTTTG-3′ #5: 5′-CCGGGCTCGCACTGACGCTTCTGTTCTCGAGAACAGAAGCGTCAGTGCGAGCTTTTT- 3′)

786-O cells were plated into 6-well plates in 2 mL of serum-supplemented RPMI 1640 medium. The day after, adherent cells were infected with COL7A1 virus (Sigma Aldrich) (1–5 MOI (multiplicity of infection)) diluted in 1 mL of serum-supplemented medium containing 8 μg/mL of polybrene (Sigma Aldrich). After 4 h, 1 mL of medium was added to each well and transduction was maintained for 16 h before changing the medium. For stable transduction, puromycin selection started 36 h post-infection (at the concentration of 2 μg/mL) and maintained for 1 month.

### Western blot

Proteins were extracted from confluent-plated cells in Laemmli buffer. Samples were heated for 5 min at 100°C. They were separated on a NuPAGE 4-12% Bis Tris gel (BioRad) at 150V for 1h15 and electro-transferred to Polyvinylidene difluoride (PVDF) membranes (BioRad) during 1h at 100V. Membranes were saturated for 1 h at room temperature in 3% BSA in TBS-Tween20 0.05% and then incubated overnight at 4°C with the appropriate primary antibody diluted in the same saturation buffer. This was followed by incubation with horseradish peroxidase (HRP)-conjugated secondary antibodies and detected by enhanced chemiluminescence. Anti-GAPDH was used as a protein loading control. Antibodies used: COL7A1 (Santa Cruz Technologies #sc33710 ; 1/1000^e^) and GAPDH (Life technologies #AM4300 ; 1/40 000^e^)

### RT-qPCR

RNA extraction was performed from confluent plated cells. Trizol was added to cell plates. After the addition of chloroform, extraction was centrifuged and the resulting upper phase, which is RNA, was precipitated with isopropanol, and the pellet was washed twice with ethanol. RNA was recovered in RNAse-free water and dosed using Nanodrop.

Reverse Transcription was achieved using the iScript cDNA synthesis kit (BioRad). Briefly, 1μg RNA was mixed with the 5x iScript Reaction Mix, Reverse Transcriptase and water. Tubes were then incubated in a thermal cycler following this cycle: 5 min at 25°C, 20 min at 46°C and 1 min at 95°C.

Real-time quantitative PCR (qPCR) was achieved thanks to the kit Prome GoTaq qPCR Master Mix (BioRad) according to manufacturer instructions. Briefly, pre-diluted (1/10) cDNA were added to the PCR plate as well as the reaction mix containing Taq polymerase and primers (10 μM) and completed with water up to a final volume of 10 μL. qPCR was run using the Biorad C1000 Thermal Cycler following this cycle: 95°C for 2 min and then, 95°C for 15 sec, 60°C for 45 sec, repeating those 2 steps 39 times and finishing by 95°C for 10 sec and 65°C for 5 sec. Primer sequences used are: #COL7A1_Fwd:5’-GGTGTTCCTACCACATGCCA-3’ ; #COL7A1_Rev:5’- GGAGGGCCGATGACTGTAAG-3’; #GAPDH_Fwd:5’- CCCATGTTCGTCATGGGTGT-3’; #GAPDH_Rev:5’-TGGTCATGAGTCCTTCCACGATA-3’; #RPL13_Fwd:5’-TTAATTCCTC ATGCGTTGCCTGCC-3’#RPL13_Rev:5’-TTCCTTGCTCCCAGCTTCCTATGT-3’. Relative quantification was performed using the comparative threshold (CT) method after determining the CT values for reference and target gene COL7A1 in each sample set according to the E−ΔΔCt method. Changes in mRNA expression level were calculated after normalization to RPL13 and GAPDH mRNA.

### Proliferation & Migration assay

Proliferation and migration were assessed using Incucyte ZOOM (Sartorius) video microscope. For proliferation, cells were plated in a 96-well plate at a density of 5000 cells per well and cell confluence was measured by taking pictures every 2 h for 72 h. Migration assay was performed by plating 30 000 cells per well in a 96-well plate. After overnight cell adhesion, a wound was made using the Wound Maker (Sartorius). A confluence of the wound over time was measured by taking pictures every 2 h for 24 h. Analysis of confluence was performed thanks to the Incucyte zoom software.

## 3. Results

### COL7A1 expression is prognostic of overall survival

With the aim of discovering novel prognostic biomarkers for ccRCC patients, we have applied a systematic approach to identify genes for which expression levels in tumor are related to overall patient survival. After the filtering processes that we described in the Material and Methods section, around 17,000 genes remained. We learned a univariable Cox model on all genes that passed filtering, and the resulting table was ordered according to the model’s P-value. The top ten genes with the lowest P-value are presented in Table 2, with COL7A1 being the one whose expression level is the most significantly linked to survival. To confirm the correlation of COL7A1 expression with survival, we learned a univariate Cox proportional hazards model on the independent E-MTAB-1980 dataset. The resulting model was significant, with a P-value of 0.0023, showing that COL7A1 is correlated to survival even in the independent dataset, with patients of different ethnic origins.

**Table 2:** Results from a univariable Cox model performed on individual genes of the TCGA:KIRC cohort. The table shows the top 10 genes with the lowest P-value. COL7A1 is found at the top of the table. HR means Hazard Ratio.

| GENE SYMBOL | P-VALUE | COEFICIENT | HR | RANK |
| --- | --- | --- | --- | --- |
| COL7A1 | 5.1x10<sup>-19</sup> | 0.44 | 1.56 | 1 |
| CDCA3 | 6.2x10<sup>-19</sup> | 0.74 | 2.09 | 2 |
| CRB3 | 2.1x10<sup>-18</sup> | -0.50 | 0.60 | 3 |
| SOWAHB | 2.5x10<sup>-18</sup> | -0.52 | 0.60 | 4 |
| SLC16A12 | 4.3x10<sup>-17</sup> | -0.28 | 0.75 | 5 |
| SORBS2 | 1.9x10<sup>-16</sup> | -0.49 | 0.61 | 6 |
| CYFIP2 | 2.3x10<sup>-16</sup> | -0.59 | 0.55 | 7 |
| METTL7A | 2.7x10<sup>-16</sup> | -0.55 | 0.57 | 8 |
| TROAP | 5.2x10<sup>-16</sup> | 0.61 | 1.83 | 9 |
| RGS17 | 1.1x10<sup>-15</sup> | 0.76 | 2.14 | 10 |

To visualize the prognostic capabilities of COL7A1, we have divided TCGA:KIRC patients into groups of High and Low expression (Figure 1a). The resulting Kaplan-Meier curve shows that patients with high COL7A1 expression in tumor have a significantly worse prognosis (log-likelihood P-value < 0.0001) exhibiting median survival of 4.3 years (3.3-5.25, 95% confidence interval (CI)) with 5-year survival probability of 43.7% (34.75-52.5%). For patients with low COL7A1 expression, it was not possible to determine median survival as the survival curve does not cross the 50% line; however, this group had a 5-year survival probability of 73.1% (67.8-78.9%).

**Figure 1:**
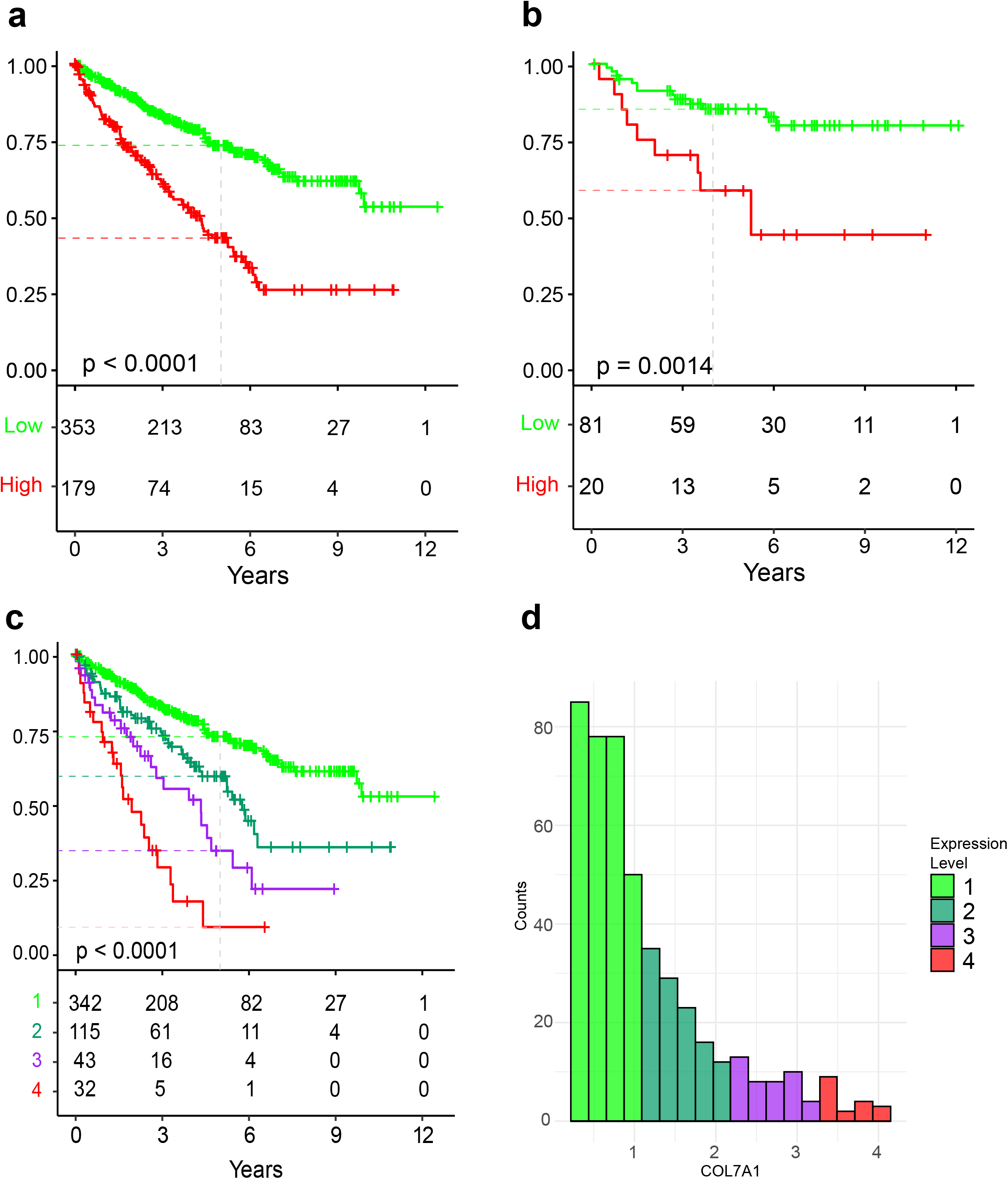
Kaplan-Meier curves showing differences in overall survival probability of patients that are grouped according to COL7A1 expression of the tumor. (a) Kaplan-Meier curve based on the TCGA:KIRC cohort where patients are grouped in groups of Low (green) and High (red) COL7A1 expression. (b) same approach as in subfigure a, applied to the E-MTAB-1980 cohort. (c) Kaplan-Meier curve based on the TCGA:KIRC cohort where patients are grouped in four groups of increasing COL7A1 expression. (d) histogram of COL7A1 expression in log2(TPM+1) where colors correspond to groups of subfigure c.

While the number of patients in the E-MTAB-1980 dataset is much lower compared to the TCGA:KIRC dataset and patients are of predominantly early stages, we can still see a similar stratification of patients according to the risk (Figure 1b). Patients in the high COL7A1 expression group have a significantly worse prognosis than the low expression group (P-value = 0.0014), confirming the results obtained on the TCGA:KIRC dataset. The 5-year survival probability of the high expression group was 58.3% (39.8-85.5%), in contrast to 85.2% (77.4-93.7%) in the low expression group.

To further investigate the prognostic power of COL7A1, we divided the range of COL7A1 expression in TCGA:KIRC dataset into four equal levels in terms of expression range (Figure 1c). The lowest level of expression is almost identical to the low expression group of Figure 1a. The other three levels of expression showed a gradually increased risk and shorter survival time of patients in relation to COL7A1 expression levels (Figure 1d). The 5-year survival probability of each group from the lowest to the highest COL7A1 expression is 72.4%, 59.1%, 34.15%, and 8.6%.

### COL7A1 expression can predict patient survival atop of clinical characteristics

Having investigated the predictive power of the standalone model based solely on COL7A1 expression, we opted to investigate how COL7A1 expression could be applied in real clinical settings. First, we investigated COL7A1 expression patterns across samples of different stages and histological grades. When compared to the overall expression of protein-coding genes in tumor samples, COL7A1 appears to be, in general, a weakly expressed gene (Figure S1a) with a median expression of 0.8 (0.74-0.856), while a median expression of all protein-coding genes is 3.058 (3.055-3.06).

Additionally, when comparing the COL7A1 tumor expression across patients with different clinicopathological characteristics, we found that COL7A1 tends to be more expressed in patients of higher stage and grade (Figure S1b and S1c). Following this finding, we investigated whether COL7A1 stratification performances remain inside these clinical characteristics (stage, grade). Indeed, we observed a significant stratification between patients with low and high COL7A1 expression in all stages (Figure 2). Stratification was most prominent in patients with stage II and IV tumor with a P-value of 0.00099 and 0.00025 respectively. The 5-year survival probability for stage-IV patients was 39.6% (95% CI 26.5%-59.2%) for the low-expression strata and 11.6% (95% CI 5.1%-26.5%) for the high-expression strata. The 5-year survival probability for stage I patients in low-expression (resp. high-expression) strata was 82.4% (resp. 69.3%), for stage II patients 87.3% (resp. 58.9%), and for stage III patients 63.5% (resp. 38.6%). We obtained similar results in two independent cohorts (Figure S2). In addition to improving prediction atop of tumor stage, COL7A1 expression was able to improve survival prediction atop of tumor grade as well (Figure S3). The most prominent effect (P-value of 0.00018) was observed in patients with Grade 3 tumors, where the 5-year survival probability of low-expression (resp. high-expression) strata was 42.7% (resp. 13.6%). Although significant, the poorest survival prediction was observed for grade 1+2 tumors, which are already of a good outcome.

**Figure 2:**
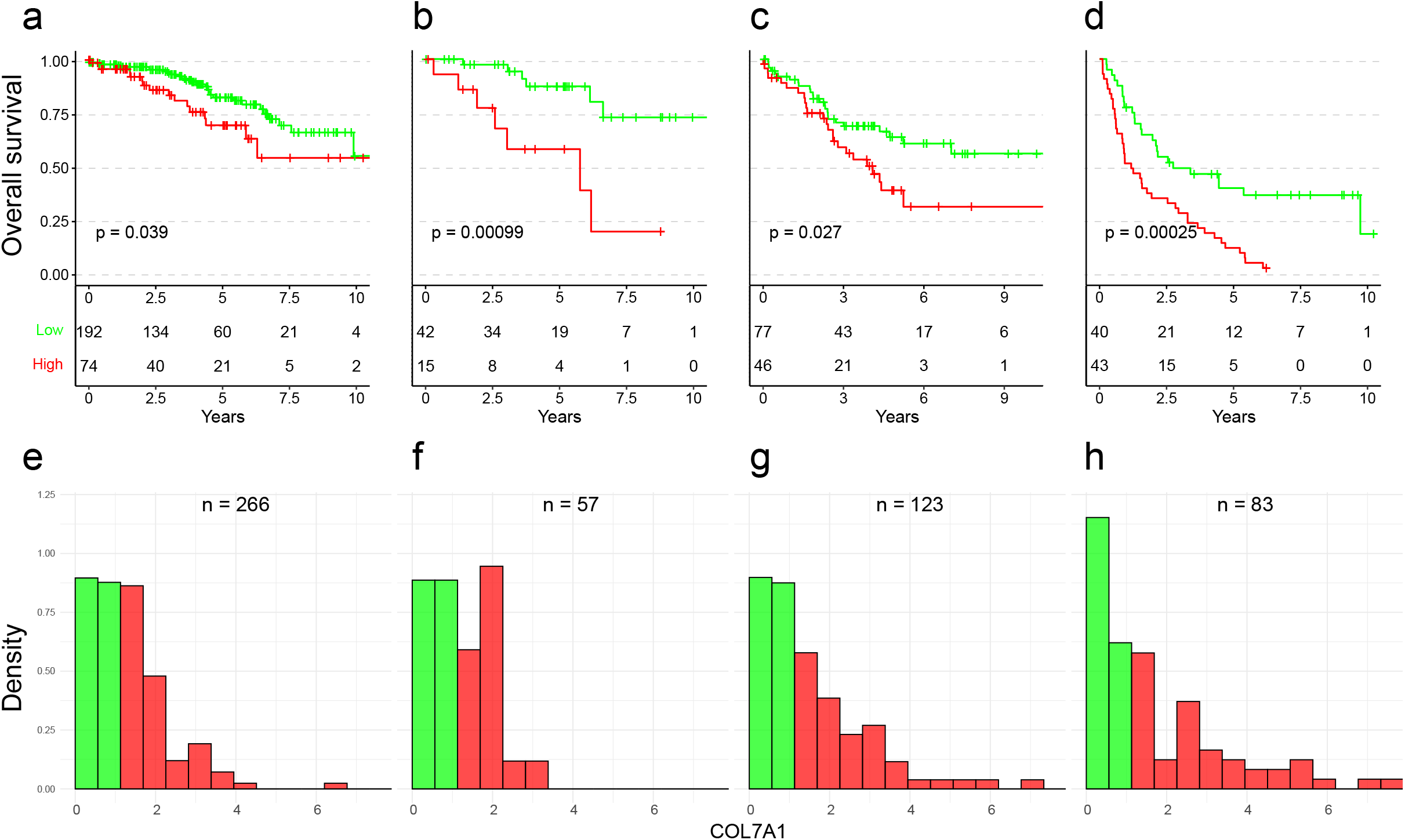
Kaplan-Meier curves and corresponding histograms showing differences in overall survival in the TCGA:KIRC cohort. Patients were split into four subsets according to the stage of patients. Patients were divided in groups of Low (green) and High (red) COL7A1 expression. (a, e) Stage I patients. (b, f) Stage II patients. (c, g) Stage III patients. (d, h) Stage IV patients. The expression threshold between the Low and High expression groups is the same as in Figure 1a.

### COL7A1 expression can improve multivariable Cox model

Using the univariable Cox model, we investigated how survival prediction based on individual clinical variables compares to prediction based on COL7A1 (Table 3). Interestingly, COL7A1 expression showed better prediction than tumor grade or age of patients. The only clinical variable that shows better prediction than COL7A1 expression is the tumor stage, especially for stage IV tumors. In a multivariable Cox model, COL7A1 keeps a particularly low P-value (P = 6.9 × 10^−9^), meaning that COL7A1 expression retains predictive power even when clinical characteristics are included in the model. To investigate what is the added value of COL7A1 expression to the multivariable Cox model based on clinical characteristics alone, we have computed an analysis of deviance table, based on the sequential partial log-likelihood. When variables were added to the model in the order they appear in the table, COL7A1 expression shows significant improvement with a P-value of 9.6 × 10^−8^ and an

**Table 3:** Results of univariable and multivariable Cox models learned on stage, grade, age of patients and COL7A1 expression in the TCGA:KIRC cohort. The table also contains results from ANOVA performed on the multivariable Cox model.

| COVARIATE | Univariable Cox model |  |  |  | Multivariable Cox model<sup>b</sup> |  |  | ANOVA |  |
| --- | --- | --- | --- | --- | --- | --- | --- | --- | --- |
| | Coefficient ($\beta$) | HR [exp($\beta$)] | P-value | Score test | Coefficient ($\beta$) | HR [exp($\beta$)] | P-value | Chisq | P-value |
| STAGE | - | - | $1.1 \times 10^{-26}$ | 123.8 | - | - | - | 95.22 | $1.7 \times 10^{-20}$ |
| II | 0.22 | 1.245 | 0.486 | - | 0.18 | 1.20 | 0.56 |  |  |
| III | 0.91 | 2.48 | $1.2 \times 10^{-05}$ | - | 0.55 | 1.73 | 0.013 | | |
| IV | 1.84 | 6.32 | $1.4 \times 10^{-21}$ | - | 1.44 | 4.23 | $1.8 \times 10^{-10}$ | | |
| GRADE<sup>a</sup> | - | - | $5.6 \times 10^{-18}$ | 79.46 | - | - | - | 14.2 | $8.3 \times 10^{-4}$ |
| III | 0.64 | 1.90 | $6.9 \times 10^{-4}$ | - | 0.36 | 1.43 | 0.067 | | |
| IV | 1.63 | 5.10 | $6.7 \times 10^{-16}$ | - | 0.60 | 1.83 | 0.011 | | |
| AGE | 0.035 | 1.03 | $2.5 \times 10^{-06}$ | 22.51 | 0.036 | 1.036 | $1.4 \times 10^{-6}$ | 22.67 | $1.9 \times 10^{-6}$ |
| COL7A1 | 0.44 | 1.56 | $5.8 \times 10^{-20}$ | 83.67 | 0.32 | 1.37 | $6.9 \times 10^{-9}$ | 28.45 | $9.6 \times 10^{-8}$ |

a) Histological grades I and II are combined due to low number of patients in grade I b) Logrank test (score) for multivariable analysis is 215.3, and Likelihood ratio test is 160.5

### COL7A1 expression is correlated with genes involved in cell division, inflammatory response and epithelial to mesenchymal transition, and anti-correlated with metabolism

To infer possible biological mechanisms, we have investigated how COL7A1 expression correlates with the expression of other protein-coding genes. As illustrated in Figure 3 **to be added ∼ here**

**Figure 3:**
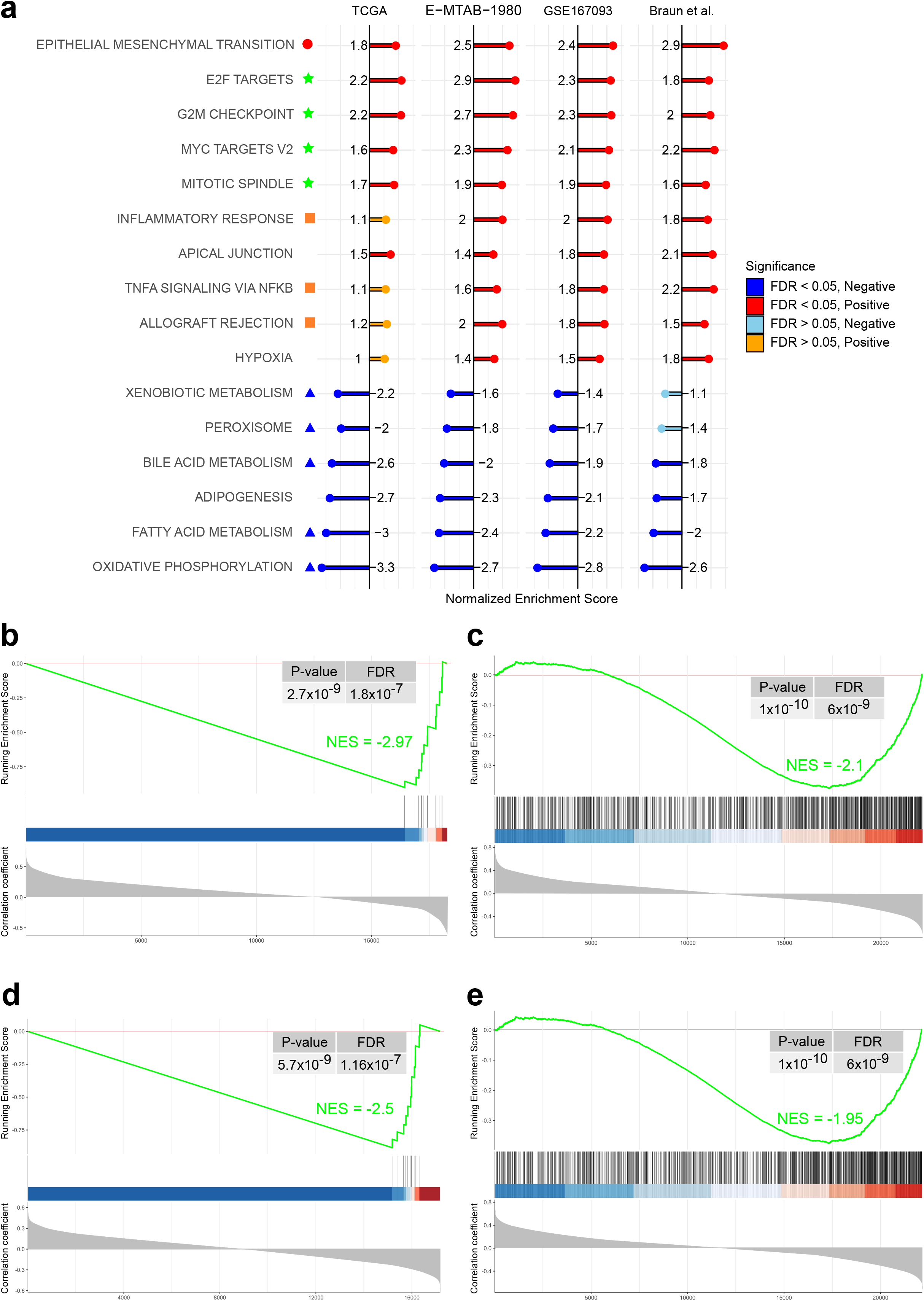
Results from GSEA analysis based on COL7A1 correlation, performed on four different datasets (TCGA:KIRC, E-MTAB-1980, GSE167093 and Braun *et al*.). (a) Most enriched HALLMARK pathways among all four datasets. Green stars highlight pathways related to cell proliferation, orange squares for inflammation/immune response, red circle for the Epithelial to Mesenchymal transition pathway, and blue triangles for metabolic pathways; (b) GSEA plot of mitochondrial genes in the TCGA:KIRC cohort; (c) GSEA plot of GO:CC MITOCHONDRION pathway in the E-MTAB-1980 cohort; (d) Same as on subfigure b, applied on the Braun *et al*. data; (e) Same as on subfigure c applied on the GSE167093 dataset.

Figure 3a, a GSEA analysis to identify the biological functions of correlated genes showed strong enrichment in cell proliferation pathways, inflammation/immune response and Epithelial to Mesenchymal transition pathway. Moreover, Supplementary Figure S4 shows that proliferation related genes are more expressed in tumors also expressing high levels of COL7A1. On the other side, the GSEA analysis showed a strong downregulation in multiple metabolic pathways with the strongest decrease being mitochondrial oxidative phosphorylation for patients with high tumor expression of COL7A1 (Figure S5). Interestingly, a strikingly reduced mitochondrial respiratory capacity has been previously observed in primary human ccRCC cells [35]. Additionally, all the 12 mitochondrial genes expressed in the TCGA dataset and Braun *et al*. showed a clear anti-correlation pattern with COL7A1 (Figure 3b and 3d and Supplementary Figure S6). Since E-TMAB-1980 and GSE167093 datasets originate from expression arrays, datasets do not contain mitochondrial genes. To investigate expression of genes for which the encoded protein is a part of mitochondria, we performed GSEA analysis on the Cellular Components (CC) of the Gene Ontology database. Most of the genes were found to be anti-correlated with COL7A1 (Figures 3c and 3e).

### Experimental validation

To validate the transcriptomic analysis, we performed *in vitro* experiments to investigate the impact of COL7A1 expression on the proliferation and migration of several RCC cell lines. First, RT-qPCR and western blot analysis revealed differential COL7A1 expression in 786-O, ACHN and Caki-1 RCC cell lines showing that the protein is abundantly expressed in the metastatic 786-O cells (Figure 4 a, b, b’). Therefore, we generated shCOL7A1 786-O cells in which we knocked down COL7A1 expression by RNA interference, using 4 different shRNAs targeting this gene. Cells infected with shRNA1 or shRNA5 COL7A1 showed a decrease in COL7A1 expression of 50 and 70% respectively (Figure 4 c, c’). Therefore, we chose these cells to further characterize them by cell proliferation and migration assays (Figure 4 d, e). After 48 h of culture, control 786-O cells were almost confluent whereas proliferation of shRNA1 or shRNA5 cells was inhibited by 10 and 20% respectively (Figure 4D and Supplementary Figure S7a). Moreover, cell migration of both shRNA cell lines assessed by the wound healing assay was significantly impaired (Figure 4e and Supplementary Figure S7b).

**Figure 4:**
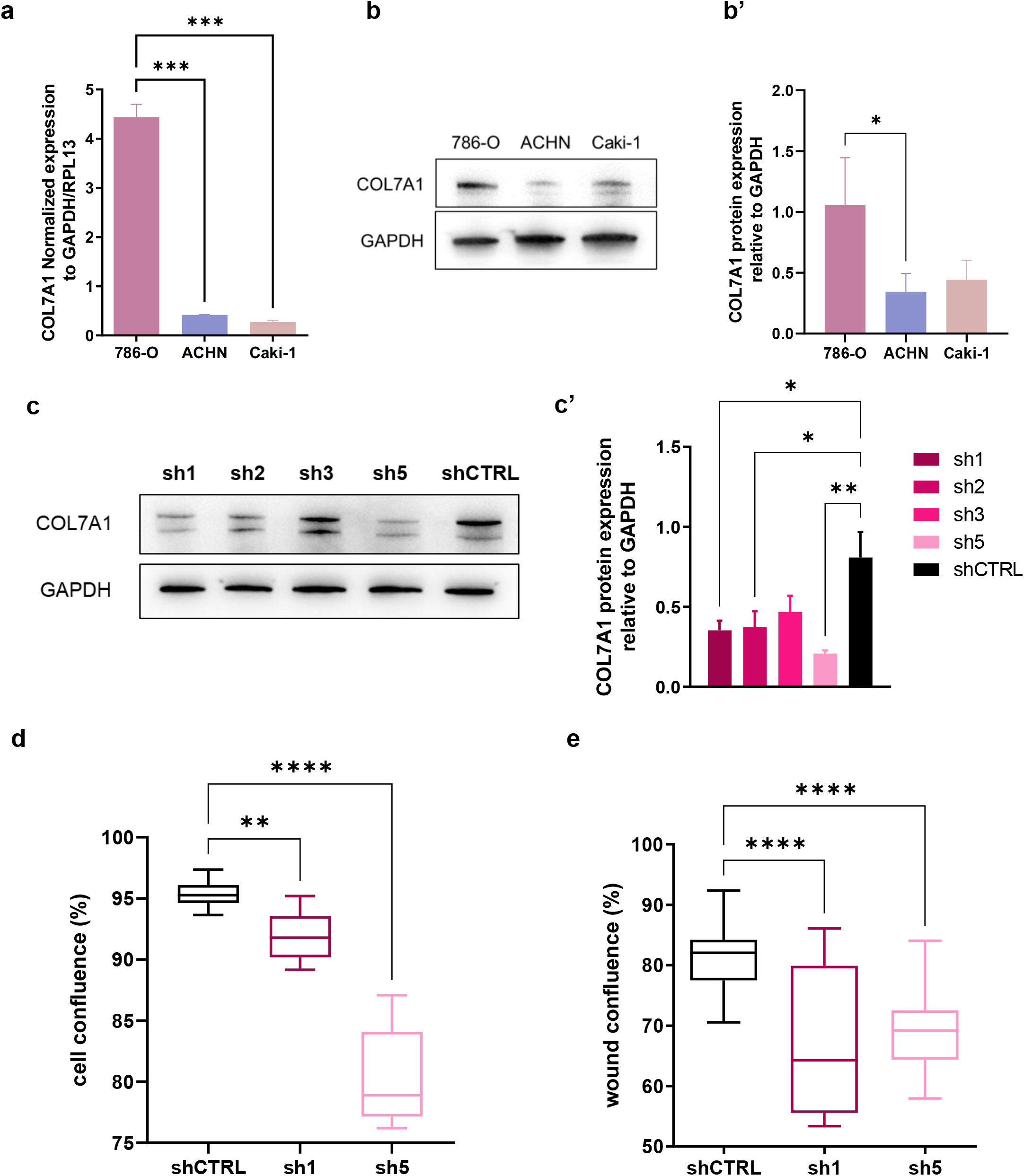
Influence of COL7A1 on proliferation and migration of 786-O ccRCC cells. (a): RT-qPCR showing expression level of COL7A1 into 3 RCC cell lines : 786-O, ACHN and Caki-1 (n=6). (b): Western blot of COL7A1 protein level into the 3 cell lines and the respective quantification relative to GAPDH (B’) (n=3). (c, c’): Western blot of shCOL7A1 786-O cell lines generated with different shRNA and its quantification (n=3). (d): Proliferation assay showing cell confluence (%) after 48 h of culturing of 786-O shCTRL, shRNA1 and shRNA5 cell lines (n= 12). (e): Wound healing assay of 786-O shCTRL, shRNA1 and shRNA5 cell lines, graph represents wound confluence (%) 10 h after the scratch (n = 24). Statistical analysis was made using unpaired t-test, p < *0.05, **0.01, ***0.001, ****0.0001.

## 4. Discussion

With the advancement of OMIC technologies and methods in machine learning, there has been huge progress in novel biomarker discovery in ccRCC [36]. In the present work, we have investigated the use of COL7A1 expression as a potential prognostic biomarker in ccRCC. Although COL7A1 is generally weakly expressed in tumor tissue (for around 70% of the patients), high expression of this gene is a sign of poor prognosis. Additionally, COL7A1 expression is inversely reciprocal to overall survival, meaning that the higher the expression of COL7A1, the poorer the patient outcome. Consequently, COL7A1 shows potential to be used either as a binary prognostic factor (low or high expression) or as a continuous prognostic factor. Future predictive models may be improved by the incorporation of new biomarkers. By stratifying the Kaplan-Meier curve according to the stage and grade of tumors, we showed that COL7A1 can improve the prediction of patient survival based on these two variables. This feature is particularly important for patients with stage II or III cancers due to their high uncertainty of outcome.

The GSEA analysis showed that high COL7A1 expression could be a sign of multiple other dysregulations in renal tumors. In particular, COL7A1 expression tends to be correlated with genes belonging to pathways that play key roles in cell proliferation, resulting in positive enrichment of “E2F targets”, “MYC targets” “G2M checkpoint” and “mitotic spindle” pathways. Alterations in these signaling pathways were observed in renal carcinoma patients [37–40], where they may lead to uncontrolled cancer cell proliferation, promoting tumor progression and aggressiveness. The GSEA analysis shows strong positive enrichment in the “Epithelial to Mesenchymal transition” pathway, a signature of dedifferentiation of normal epithelial renal cells, as previously reported in a study comparing ccRCC cells to tumor-adjacent kidney tissue [4]. Those results are further supported by our *in vitro* assays showing that proliferation and migration of 786-O cells are both impacted upon COL7A1 knockdown.

We also observe anti-correlation between COL7A1 and mitochondrial genes, as well as genes involved in various metabolic pathways. Warburg effect is known to be prominent in ccRCC and is known to play a role in epigenetic changes of cancer cells [41].

Thus, the incorporation of COL7A1 expression as a valid biomarker into increasingly personalized predictive tools may help to predict outcomes for patients with renal cell carcinoma. Further research on metabolic deregulations in ccRCC as well as functional implications of COL7A1 regulations are needed.

## Author Contributions

Conceptualization, L.G. and O.F.; data analysis, D.K., L.G. and R.J.; experiments, I.S. and O.F.; original draft preparation, D.K, C.C., I.S., O.F. All authors have reviewed and accepted the manuscript.

## Funding

This research was supported in part by University Grenoble Alpes, Commissariat à l’Energie Atomique et aux Energies Alternatives (CEA), Institut National de la Santé et de la Recherche Médicale (Inserm). This article was developed in the framework of the Grenoble Graduate School in Chemistry, Biology and Health CBH-EUR-GS (ANR-17-EURE-0003), and Labex GRAL (ANR-10-LABX-49-01).

## Acknowledgments

The authors would like to thank Kilian Laho for his technical support. The results <published or shown> here are in whole or part based upon data generated by the TCGA Research Network: https://www.cancer.gov/tcga

Institutional Review Board Statement: Not applicable.

## Informed Consent Statement

This study uses publicly available human datasets. The consent from patients are gathered with the original studies.

## Data Availability Statement

Publicly available datasets were analyzed in this study. These data can be found here: https://www.cancer.gov/tcga ; https://www.ebi.ac.uk/arrayexpress/experiments/E-MTAB-1980/ ; supplementary data of [26] ; https://www.ncbi.nlm.nih.gov/geo/query/acc.cgi?acc=GSE167093 ;. The R script to reproduce the results presented in this article is available at **https://github.com/IRIG-BCI-IMAC/COL7A1-project** (all accessed on 19 December 2022).

## Conflicts of Interest

The authors declare no conflict of interest.

## Notes

### Competing Interest Statement

The authors have declared no competing interest.

