## Supplementary Figures for "COL7A1 expression improves prognosis prediction for patients with clear cell renal cell carcinoma atop of stage"

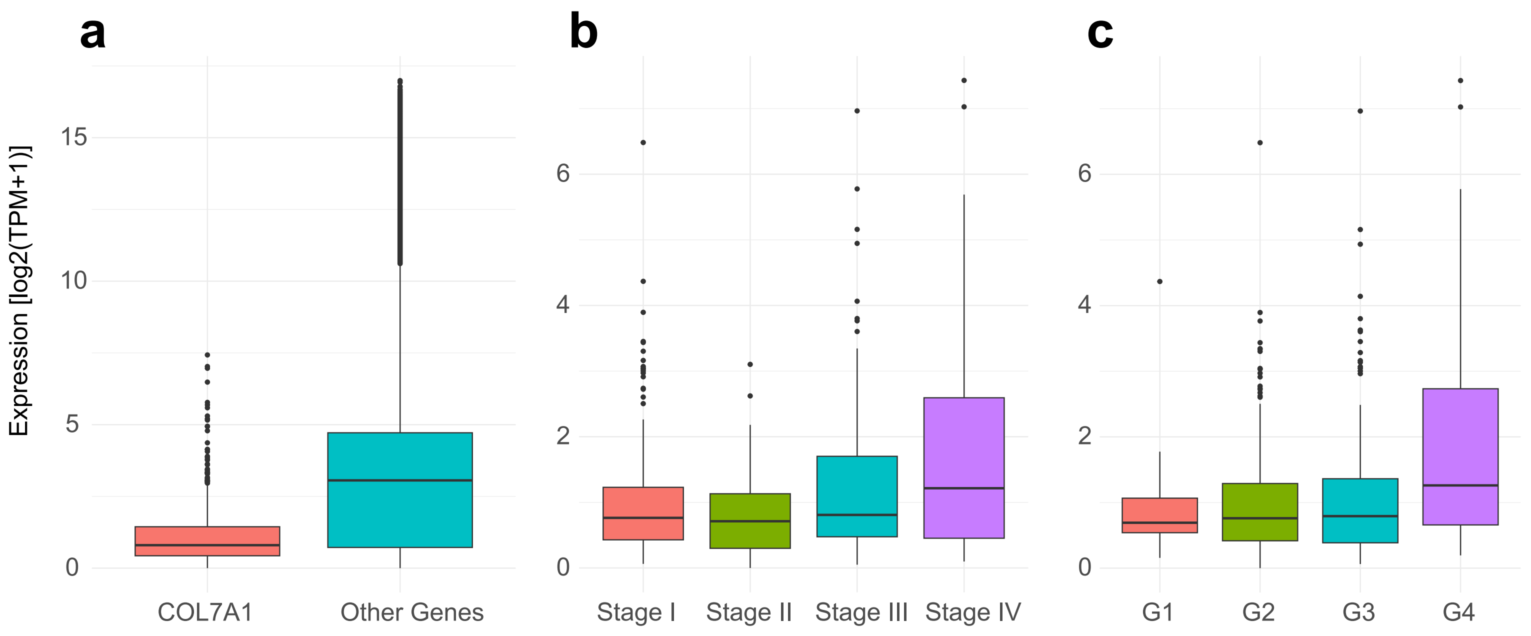


**Supplementary Figure S1:** patterns of COL7A1 expression. Expression is in log2(TMP+1) : (**a**) boxplot comparing the expression of COL7A1 and all other expressed protein-coding genes in the TCGA:KIRC cohort. Genes that are expressed less than 10 TPM across all samples are not shown on the plot. (**b**) boxplot comparing COL7A1 expression among different tumor stages; (**c**) boxplot comparing COL7A1 expression among different histological grade tumors.

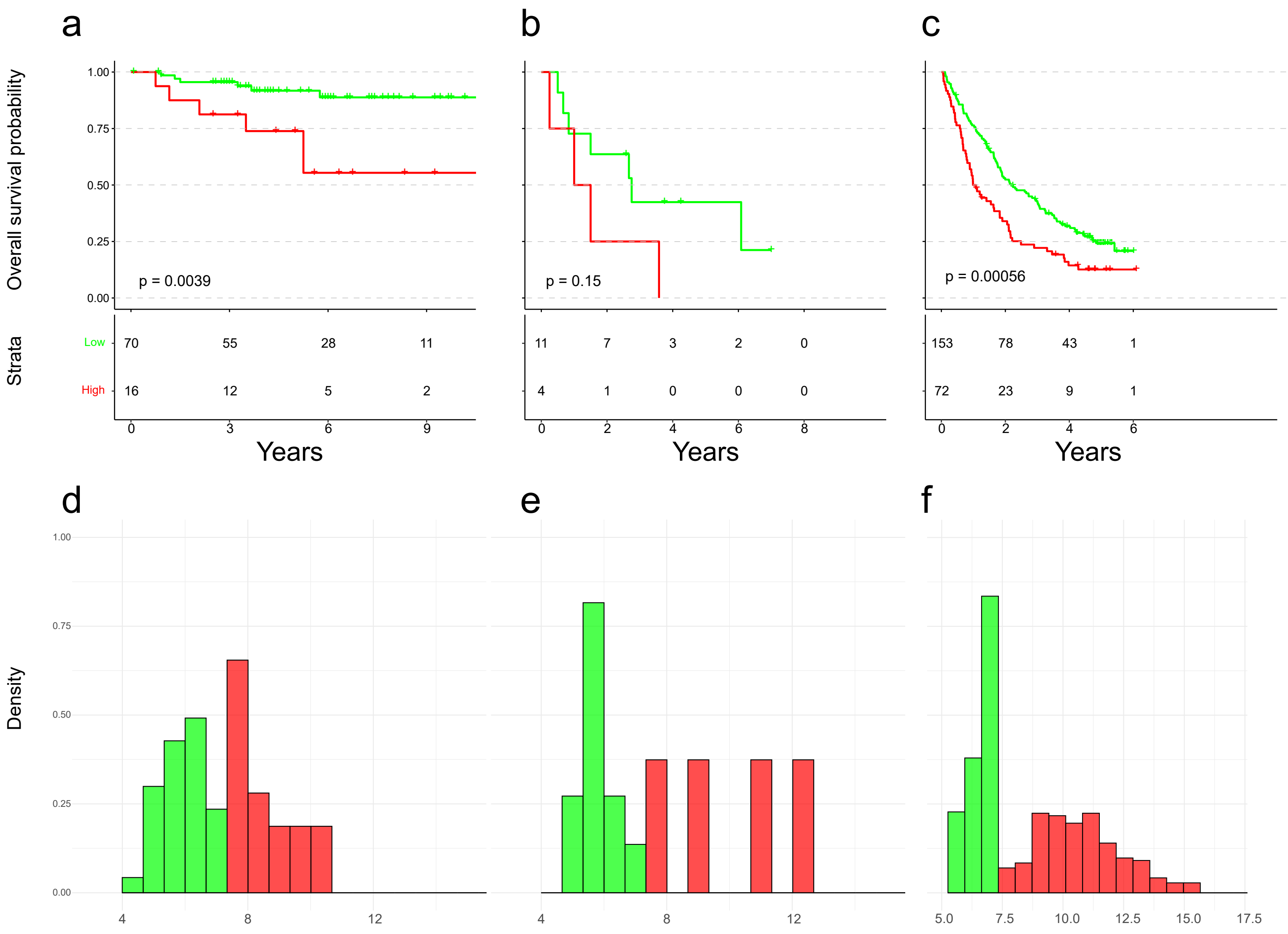


Supplementary Figure S2: Kaplan-Meier curves and corresponding histograms showing differences in overall survival in the E-MTAB-1980 and the Braun et al. cohorts. Patients were divided into groups of Low (green) and High (red) COL7A1 expression. The expression threshold between Low and High expression groups is the same as for Figure 1b. Patients of the E-MTAB-1980 cohort were split in two groups according to the stage. (**a,d**) stages I and II. (**b,e)** stages III and IV. (**c,f**) Braun et al. cohort, containing only stage IV patients.

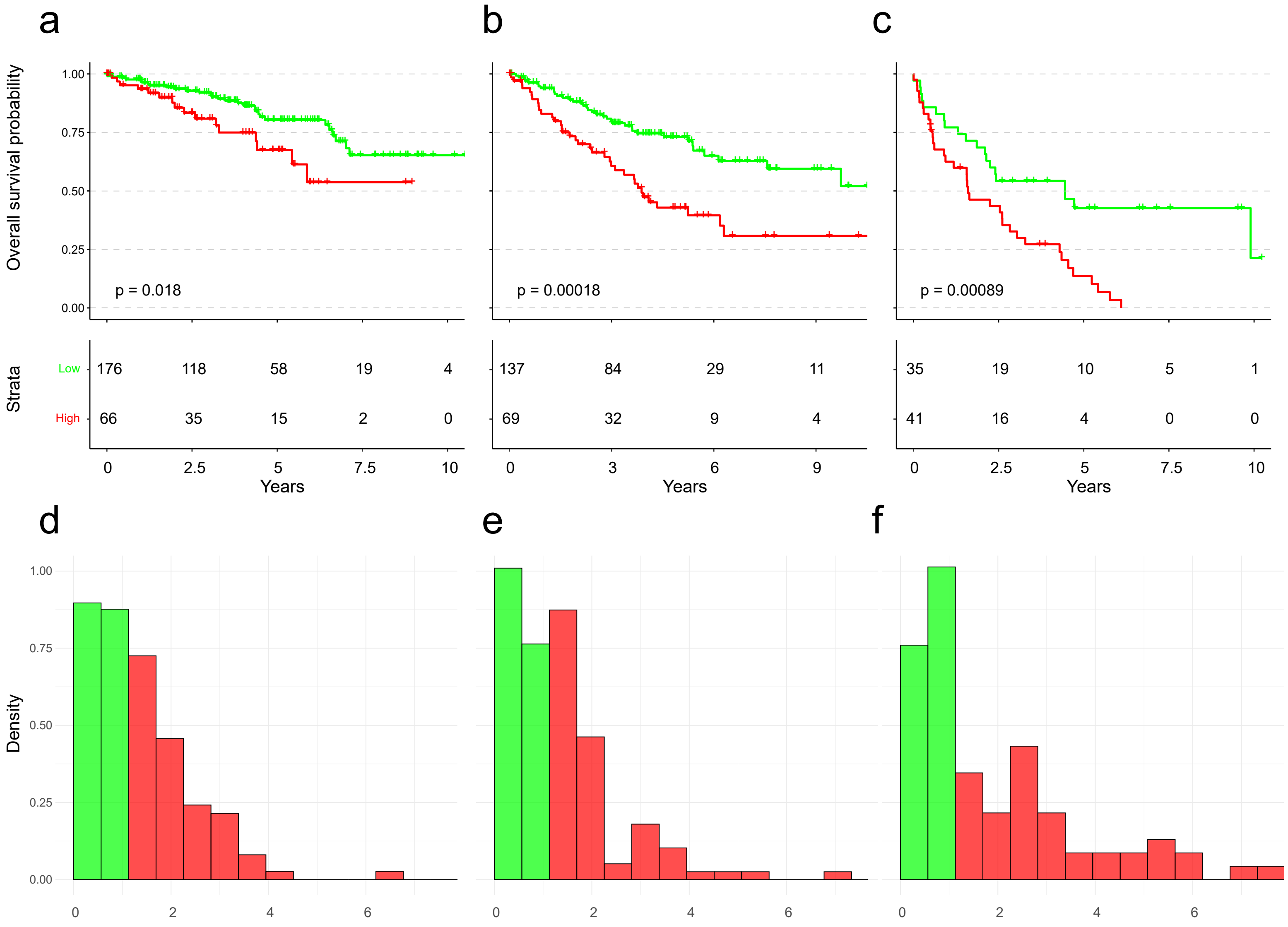


Supplementary Figure S3: Kaplan-Meier curves and corresponding histograms showing differences in overall survival of the TCGA:KIRC cohort. Patients were split in three subsets according to the grade of patients. Patients were divided in groups of Low (green) and High (red) COL7A1 expression. Expression threshold between Low and High expression groups is the same as for Figure 1A. (a,d) Grade I and II patients. (b,e) Grade III patients. (c,f) Grade IV patients.

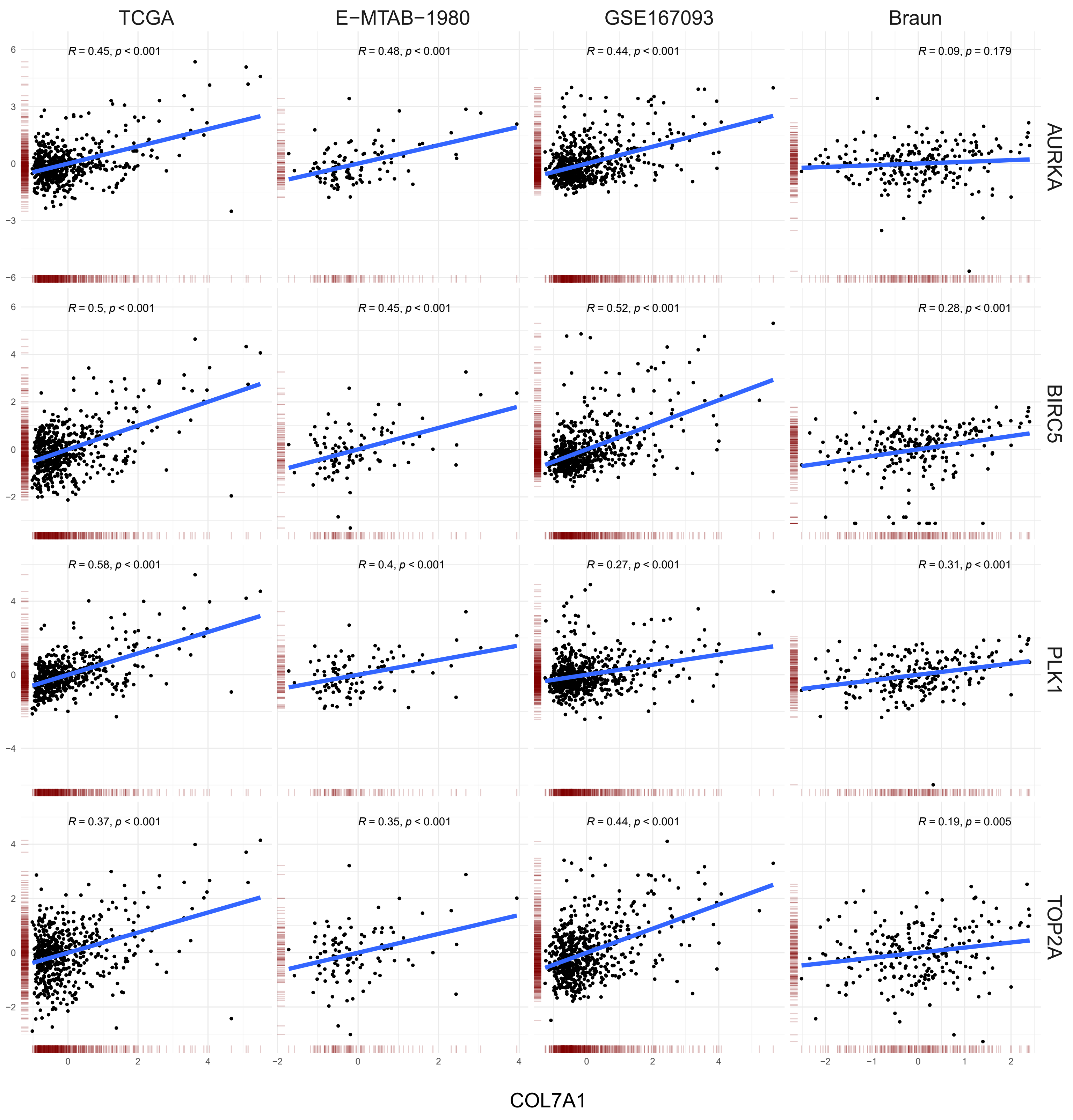


Supplementary Figure S4: Graphical representation of correlation between COL7A1 (x-axis) and four genes that belong to proliferation-related pathways (y-axis). Gene expression is scaled for better comparability.

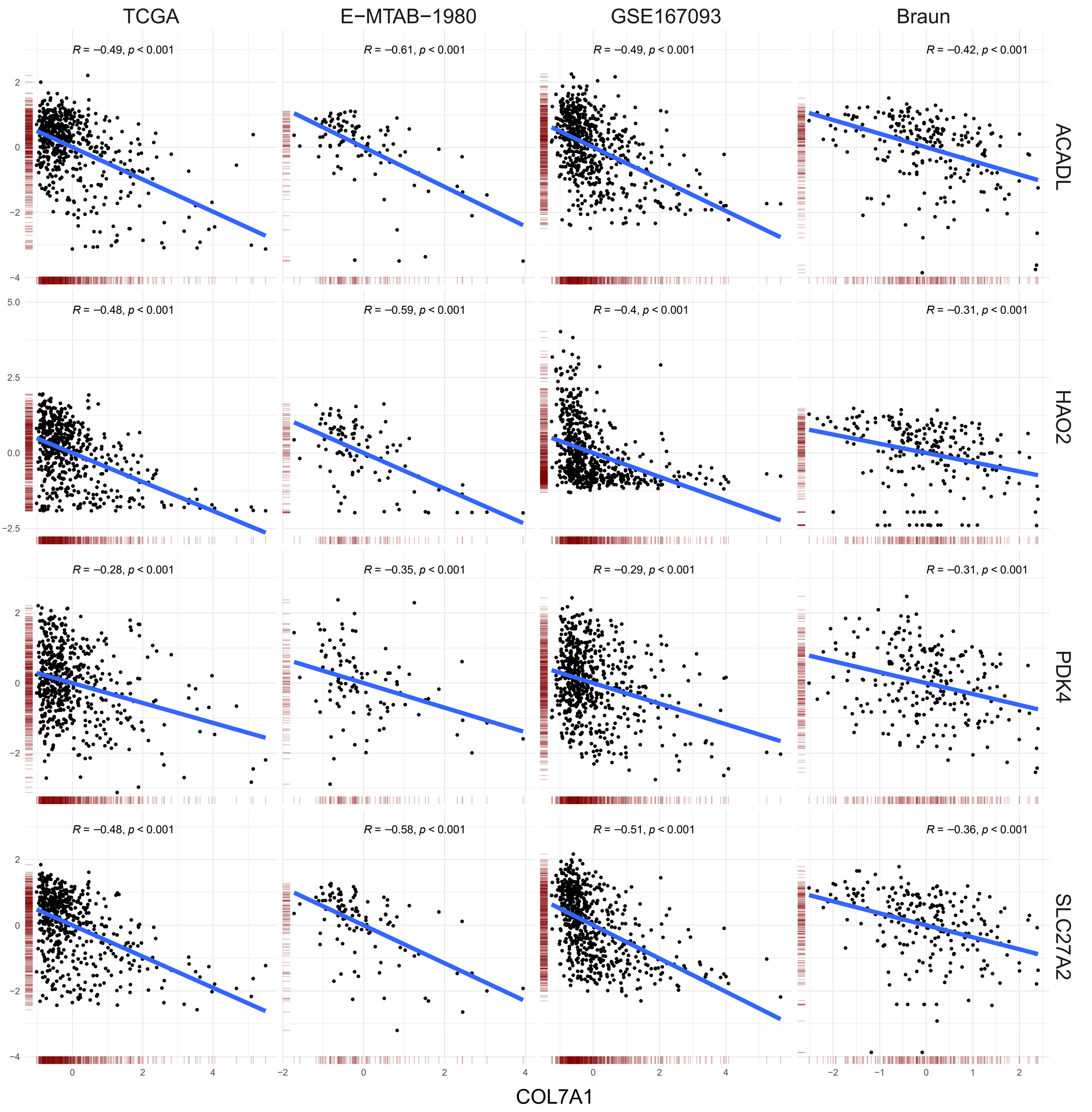


Supplementary Figure S5: Graphical representation of correlation between COL7A1 (x-axis) and four genes that belong to metabolism-related pathways (y-axis). Gene expression is scaled for better comparability.

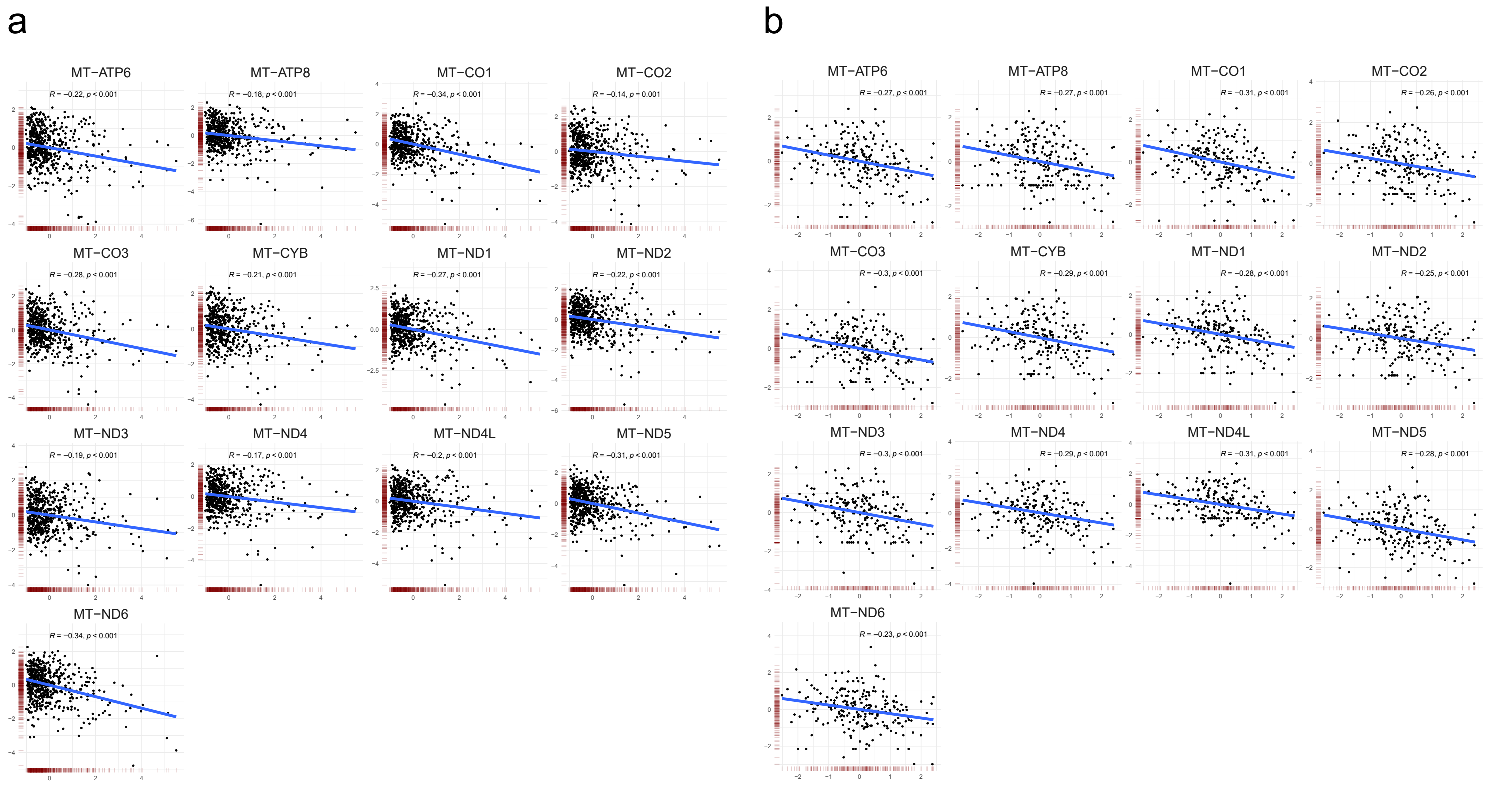


Supplementary Figure S6: Graphical representation of correlation between COL7A1 (x-axis) and mitochondrial genes (y-axis). Gene expression is scaled for better comparability. (a) TCGA:KIRC dataset; (b) Braun *et al.* dataset.

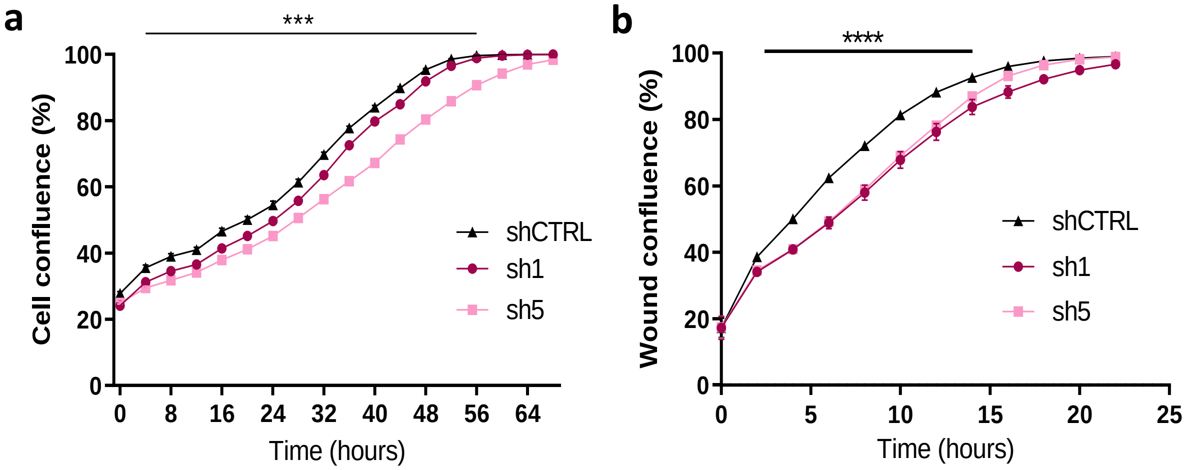


Supplementary Figure S7 Proliferation and Migration assay kinetics of 786-O shCOL7A1 cell lines. (a) Proliferation assay showing confluence of cells over time for 70 hours (n=12). (b) Scratch assay showing confluence of the wound over time for 24 hours (n=24). Statistical analysis was made using Two-Way ANOVA $p <$ *0.05, **0.01, ***0.001, ****0.0001.
